# THE COMPLEX HODOLOGICAL ARCHITECTURE OF THE MACAQUE DORSAL INTRAPARIETAL AREAS AS EMERGING FROM NEURAL TRACERS AND DW-MRI TRACTOGRAPHY

**DOI:** 10.1101/2021.02.16.431444

**Authors:** Roberto Caminiti, Gabriel Girard, Alexandra Battaglia-Mayer, Elena Borra, Andrea Schito, Giorgio M. Innocenti, Giuseppe Luppino

## Abstract

In macaque monkeys, dorsal intraparietal areas are involved in several daily visuo-motor actions. However, their border and sources of cortical afferents remain loosely defined. Through a retrograde tracer and MRI diffusion-based tractography study here we show a complex organization of the dorsal bank of the IPS, which can be subdivided into a rostral area PEip, projecting to the spinal cord, and a caudal area MIP lacking such projections. Both areas include a rostral and a caudal sector, emerging from their ipsilateral, gradient-like connectivity profiles. As tractography estimations, we used the cross-sectional volume of the white matter bundles connecting each area with other parietal and frontal regions, after selecting ROIs corresponding to the injection sites of retrograde tracers. A quantitative analysis between the proportions of cells projecting to all sectors of PEip and MIP along the continuum of the dorsal bank of the IPS and tractography revealed a significant correlation between the two data sets for most connections. Moreover, tractography revealed “false positive” but plausible streamlines awaiting histological validation.

## INTRODUCTION

Areas PEip and MIP in the dorsal bank of the intraparietal sulcus (db-IPS) of monkeys are two crucial nodes for controlling visuomotor behavior. This view stems from different sources of information. The first relates to their input-output relationships (Johnson et al., 1996; Caminiti et al., 1996; Matelli et al., 1998; Marconi et al, 2001; Bakola et al., 2017; Battaglia-Mayer and Caminiti, 2019), since they receive projections from visuomotor areas V6A and PGm and project to premotor and motor cortex (see Caminiti et al., 2017). The second consists in the functional properties of their neurons (see Lacquaniti et al., 1995; Johnson et al., 1996; Batista et al., 1999), most of which combine retinal signals about target location, with eye and hand position and movement direction signals within their tuning fields (Battaglia-Mayer et al., 2000, 2001). The third stems from the analysis of the consequences of lesions of the putative homologue areas in humans, consisting in a defective visual control of reaching, known as optic ataxia (Bálint, 1909; see Rossetti and Pisella, 2018).

To date, aspects of PEip and MIP connectivity remain obscure, since uncertain is their border. In fact, injections of retrograde and/or anterograde tracers can hardly fill the cortex of the entire dorso-ventral extent of the IPS, rendering only a partial view of its connectivity. Previous attempts to mark this border were based on the presence of cortico-spinal projections in PEip vs. MIP (Matelli et al., 1998) or on myeloarchitectonic criteria (Bakola et al., 2017). To date, no study has provided a parcellation of the cortex of the db-IPS based on cytoarchitectonics, beyond Pandya and Seltzer (1982), who labelled this large region of the superior parietal lobule (SPL) as area PEa. This study was, however, antecedent to the identification of MIP as the dorsal intraparietal region projecting to area PO (Colby et al., 1988).

The difficulties of histological studies can tentatively be overcome by diffusion-weighted MRI tractography (DW-MRI). Albeit known limitations, such as the identification of false-positive connections and biases toward reconstructing short and strong connections (Jones et al., 2013; Van Essen et al., 2014; Jbabdi et al., 2015; Knosche et al., 2015; Jeurissen et al., 2017; Maier-Hein et al., 2017; Aydogan et al., 2018; Schilling et al., 2019a,b; Girard et al., 2020), tractography shows promising results when compared to histology (Dauguet et al., 2007; Dyrby et al., 2007; Seehaus et al., 2012; Jbabdi et al., 2013; Thomas et al., 2014; Azadbakht et al., 2015; Calabrese et al., 2015; Gyengesi et al., 2015; van den Heuvel et al., 2015; Knosche et al., 2015; Donahue et al., 2016; Delettre et al., 2019; Ambrosen et al., 2020; Girard et al., 2020). Particularly, Calabrese et al. (2015), Donahue et al. (2016) and Ambrosen et al. (2020) have reported positive results when comparing labelled cells count from tracer injections in the monkey brain with connectivity weights derived from DW-MRI tractography.

In this study, we combined DW-MRI tractography and histology to elucidate the connectivity of PEip and MIP. In two macaque monkeys, we injected different retrograde fluorescent tracers along the A-P extent of the db-IPS and established their putative border based on the distribution of cortico-spinal cells projecting to the cervical segments of the spinal cord, as determined in two other animals (see Matelli et al., 1998). The connectivity of the db-IPS was studied with tractography in a fifth animal and compared in a quantitative fashion with histological data. To explore potential connections of PEip and MIP not yet revealed by tract tracing studies, the dorso-ventral extent of these areas was subdivided into different regions of interest (ROIs). This was based on the results of earlier anatomo-functional studies (Johnson et al., 1996; Battaglia-Mayer et al., 2001) showing systematic changes of both functional properties and cortico-cortical connectivity in the dorso-ventral extent of the intraparietal cortex.

Combining histology and tractography revealed a promising correlation between the proportion of cells projecting to MIP and/or PEip and the diffusion-based connectivity estimates of the corresponding streamlines revealed by tractography. Beyond advancing the information about the connectivity of the IPS, these results offer a quantitative cross-validation of the two methods within network suitable for rigorous quantitative analysis and call for a histological validation of predictions emerging from tractography.

## RESULTS

### Neural Tracers Study

#### Subdivision of the db-IPS and location of the injection sites

The location of the injection sites placed at different A-P levels in the db-IPS and involving the bank for several mm in depth (cases 72 and 73) is shown in Figure 1. To assign injection sites and RLC in the db-IPS to specific cortical entities, as in Matelli et al. (1998), we subdivided this region based on the distribution of corticospinal neurons, which clearly distinguishes between a rostral and a caudal sector (Fig. 2).

**Figure 1.**
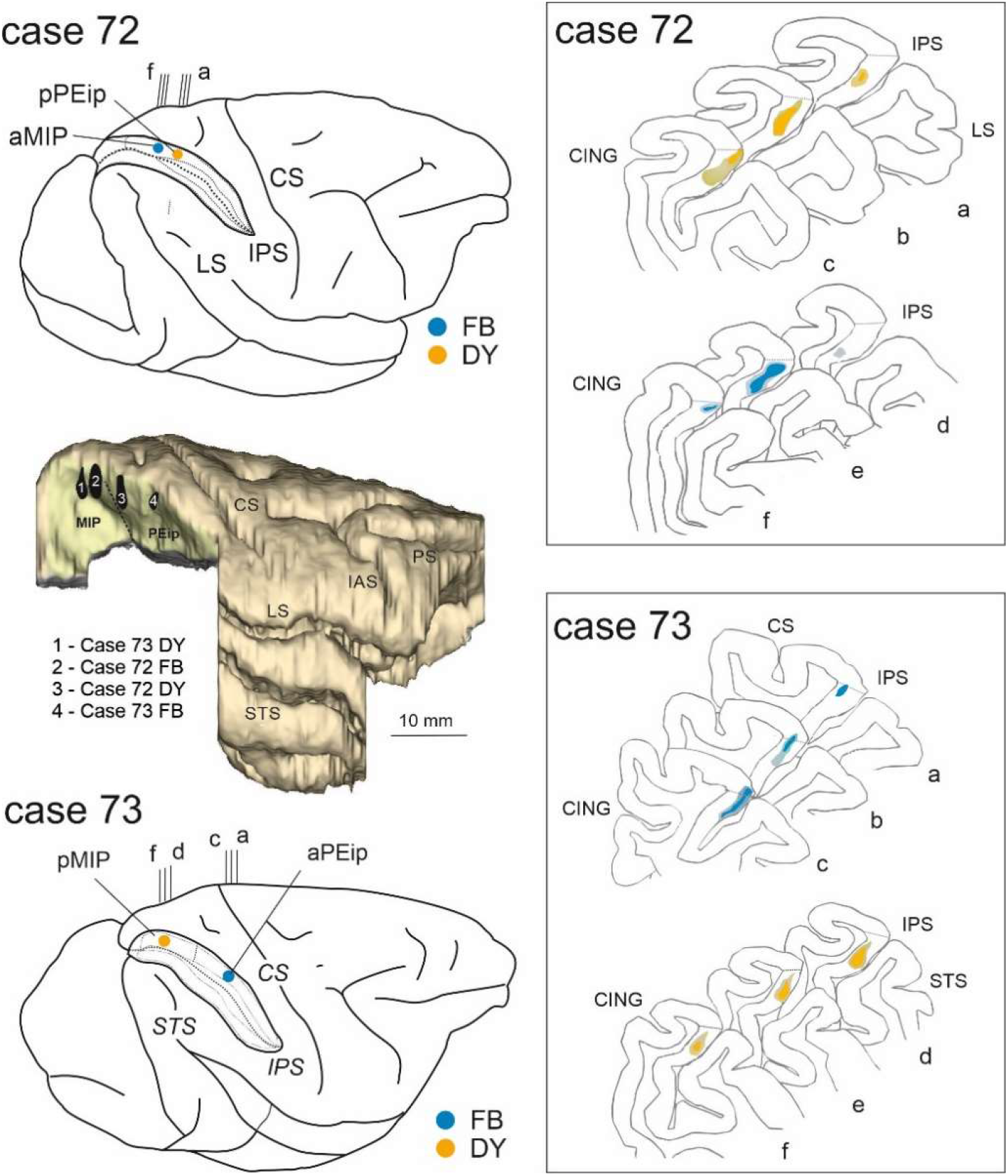
Top and bottom panels show the brain figurines with injections sites (left) of DY and FB along the db-IPS (IPS) and the corresponding histological sections (right) for cases 72 and 73. The IPS is shown as “opened” to better visualize the dorsal and ventral banks. pPEip and aPEip indicate anterior and posterior part of area PEip, respectively. The same applies to area MIP (aMIP, pMIP). In the section drawings, the injection sites are shown as a deep colored zone corresponding to the core surrounded by a light-colored zone corresponding to the halo. The middle panel on the left is a 3-D reconstruction of part of the macaque monkey left hemisphere in which the inferior parietal lobule, including the ventral bank of the IPS, was removed to show in a single comprehensive image the relative antero-posterior locations of the four tracer injections (black spots, 1-4) in the different sectors of areas MIP and PEip. CS, STS, LS, PS, lAS, and CING indicate central, superior temporal, lateral, principal, arcuate (lateral limb) and cingulate sulci.

**Figure 2.**
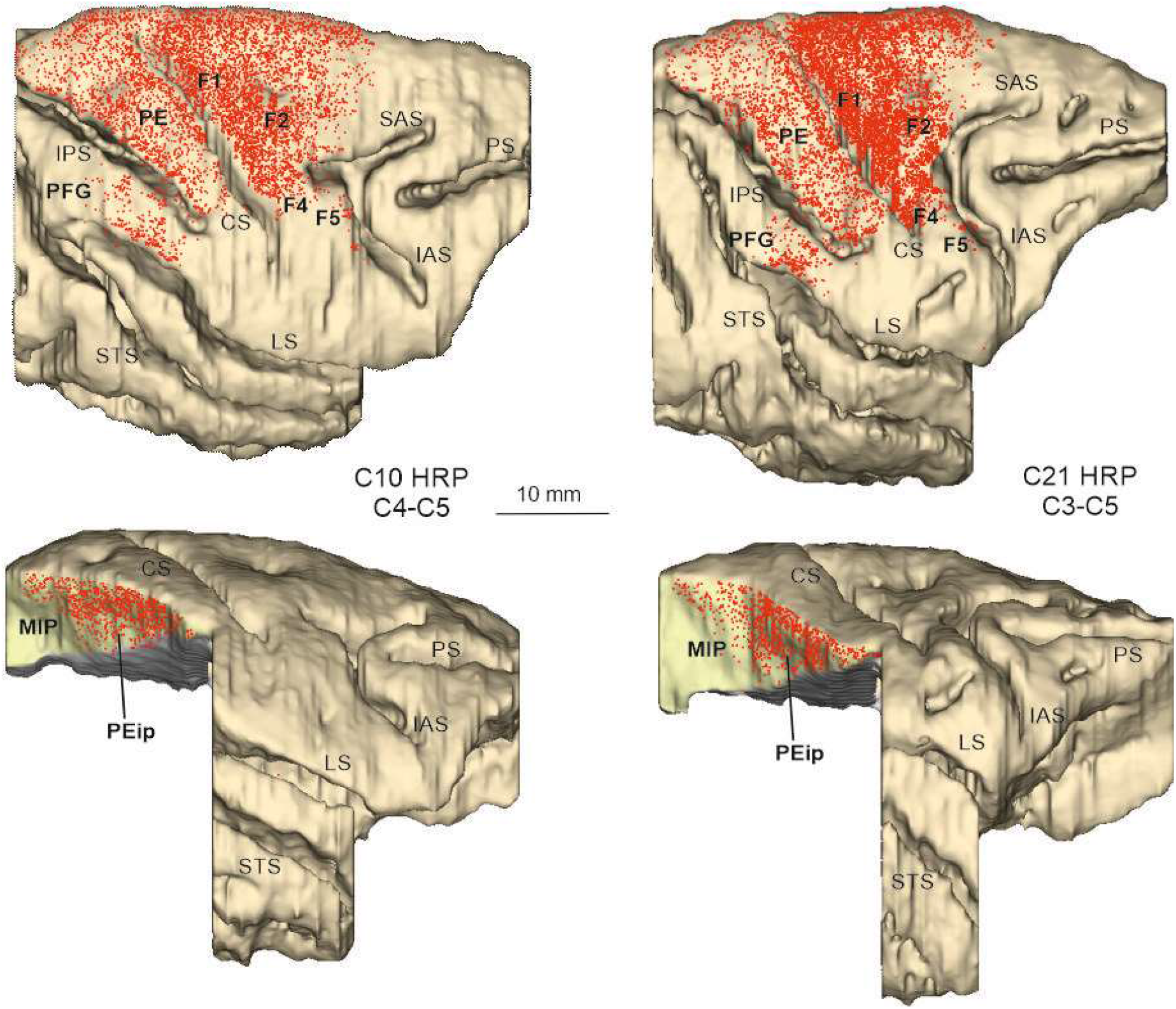
Distribution of RLC observed following HRP injections in the lateral funiculus of the spinal cord at upper cervical levels in Cases 10 and 21, shown in dorsolateral views of the 3D reconstructions of the injected hemispheres and lateral views of the db-IPS exposed after dissections of the inferior parietal lobule and of part of temporal lobe. Each dot corresponds to one labelled neuron. SAS = superior arcuate sulcus. Other abbreviations as in Figure 1.

The upper part of Figure 2 shows the overall distribution on the dorsolateral cortical surface of the corticospinal labelled neurons observed after the injection of HRP in the lateral funiculus at the upper cervical levels (Cases 10 and 21). The extensive labelling observed in both cases all over the precentral and postcentral gyri, except their most lateral part, suggested complete, or almost complete involvement of the contralateral lateral funiculus by the HRP injection. In the lower part of Figure 1, lateral views of the two hemispheres show the distribution of the RLC observed in the db-IPS. In both hemispheres, the rostral part of the bank hosted the highest number of them, as compared to its caudal part, from the crown to the fundus. This rostral sector, which does not appear to project to the thoraco-lumbar spinal cord (Matelli et al., 1998) and hosts neurons dysinaptically connected with hand motorneurons (Rathelot et al., 2017), has been here referred to as to PEip, according to the original definition of Matelli et al. (1998). Caudal to PEip, corticospinal neurons appeared to be confined to the uppermost part of the bank, which, therefore, for most of its extent lacked these projections. This last sector as a whole has been here referred to as area MIP. The border between PEip and MIP tended to run obliquely, from a ventro-rostral to a dorso-caudal position and, at about half of the depth of the bank appeared to be located at an A-P level of about 13 mm caudal to the rostral end of the IPS. In the caudalmost part of the bank, MIP borders caudally with V6A (Luppino et al., 2005; Bakola et al., 2017).

#### Ipsilateral cortical projections to area MIP

Two tracer injections targeted MIP (Fig.1), one in Case 72, where DY was placed in aMIP and one in Case 73, where FB was delivered in pMIP. The analysis of the distribution of RLC in the ipsilateral hemisphere revealed substantial labelling in both frontal and parietal areas with a smaller contribution from selected cingulate zones (Table 1). The results from these two injections will be described together and are illustrated in Figures 3–5.

**Table 1.**
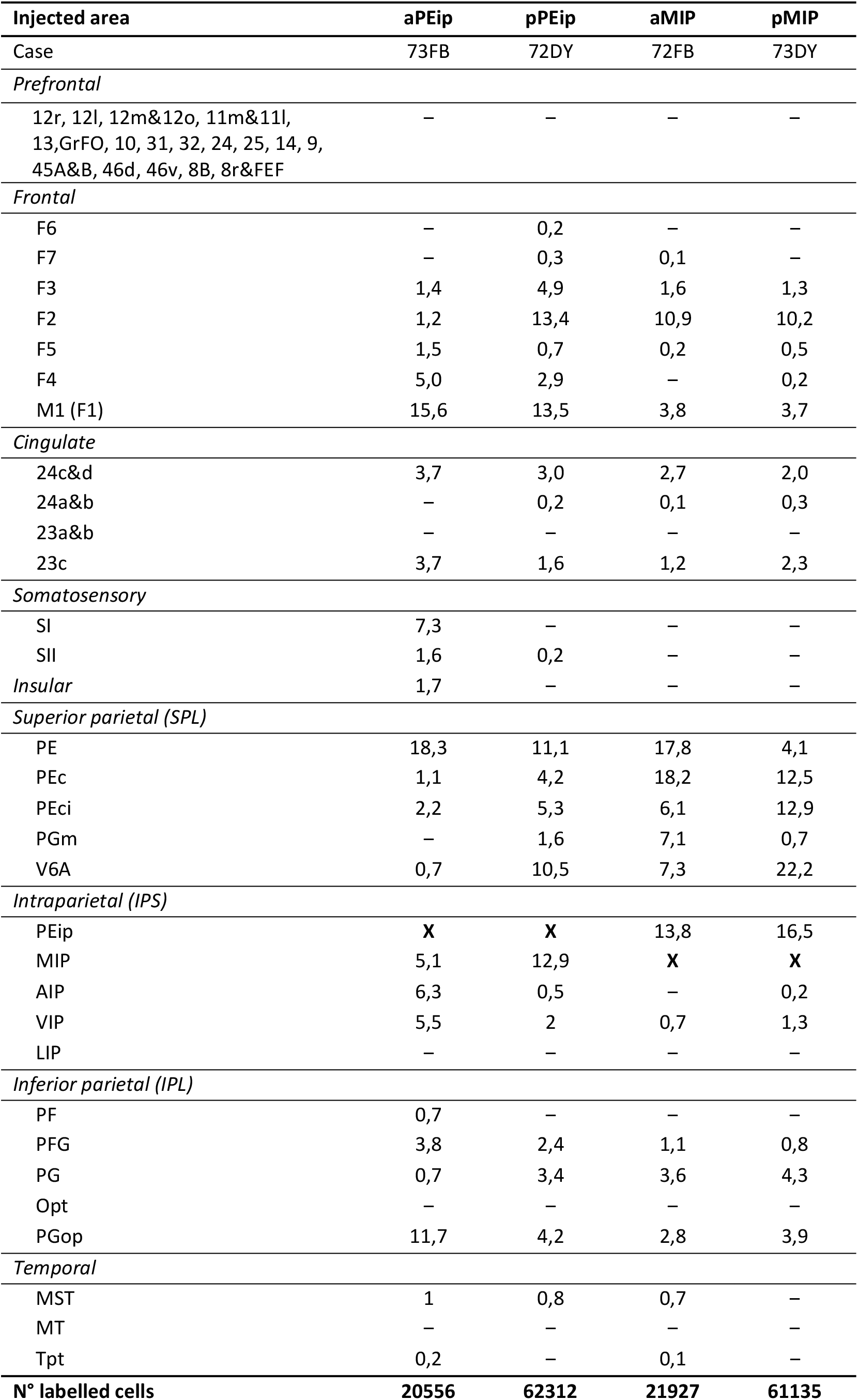
Distribution (%) and total number (n) of labelled neurons observed after tracer injections in MIP and PEip. Injection sites are sorted relative to their antero-posterior position along the db-IPS, to better display the gradient-like distribution of their projections (–, labelling < 0,1 % or no labelling). No cell counts are reported for the areas containing the injection sites (**X**).

**Figure 3.**
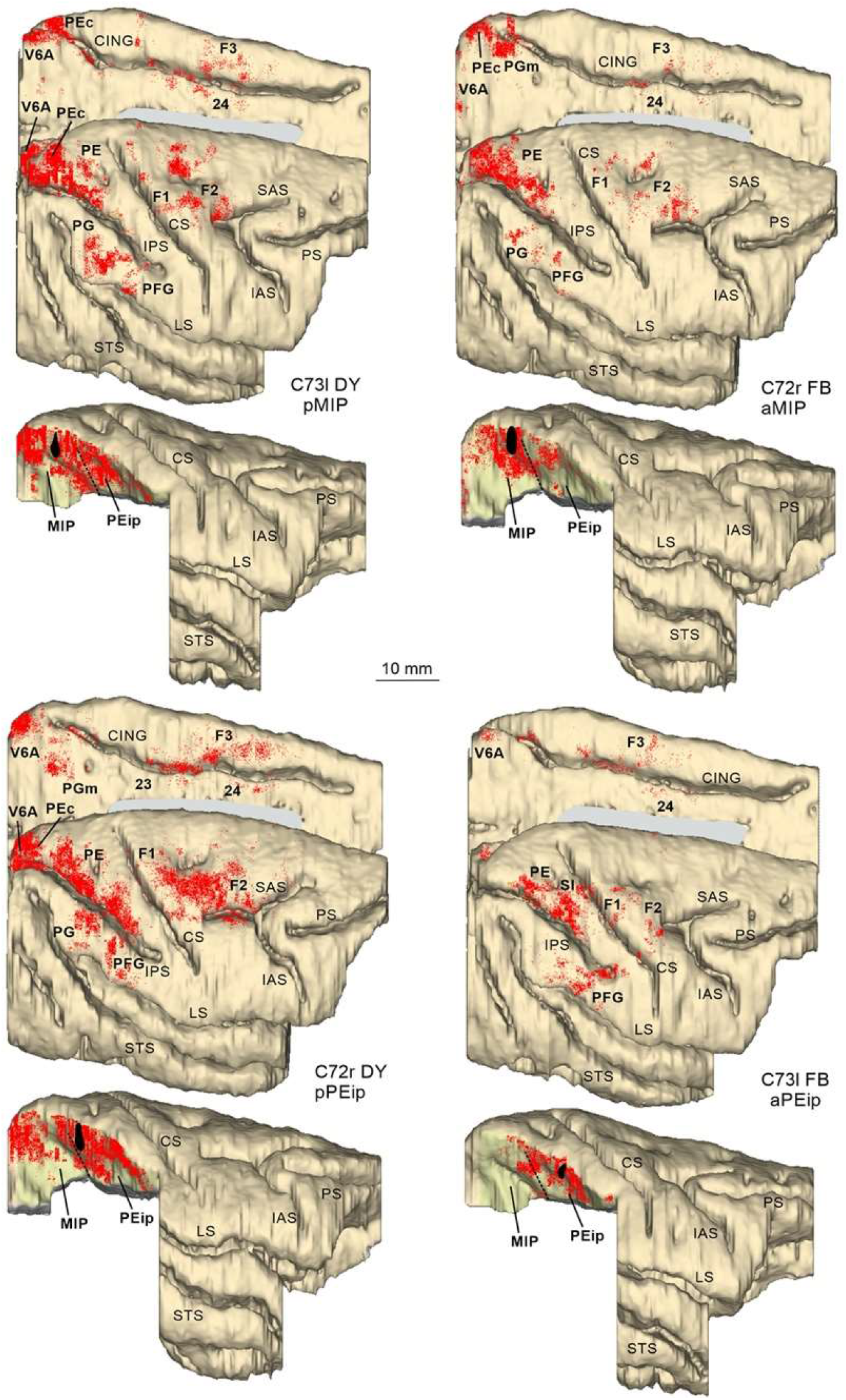
Distribution of RLC observed following tracer injections in the db-IPS, shown in dorsolateral and mesial views of the injected hemispheres and in lateral views of the db-IPS. The hemisphere of Case 73 is shown as a right hemisphere. Abbreviations and conventions as in Figures 1 and 2.

**Fig. 4.**
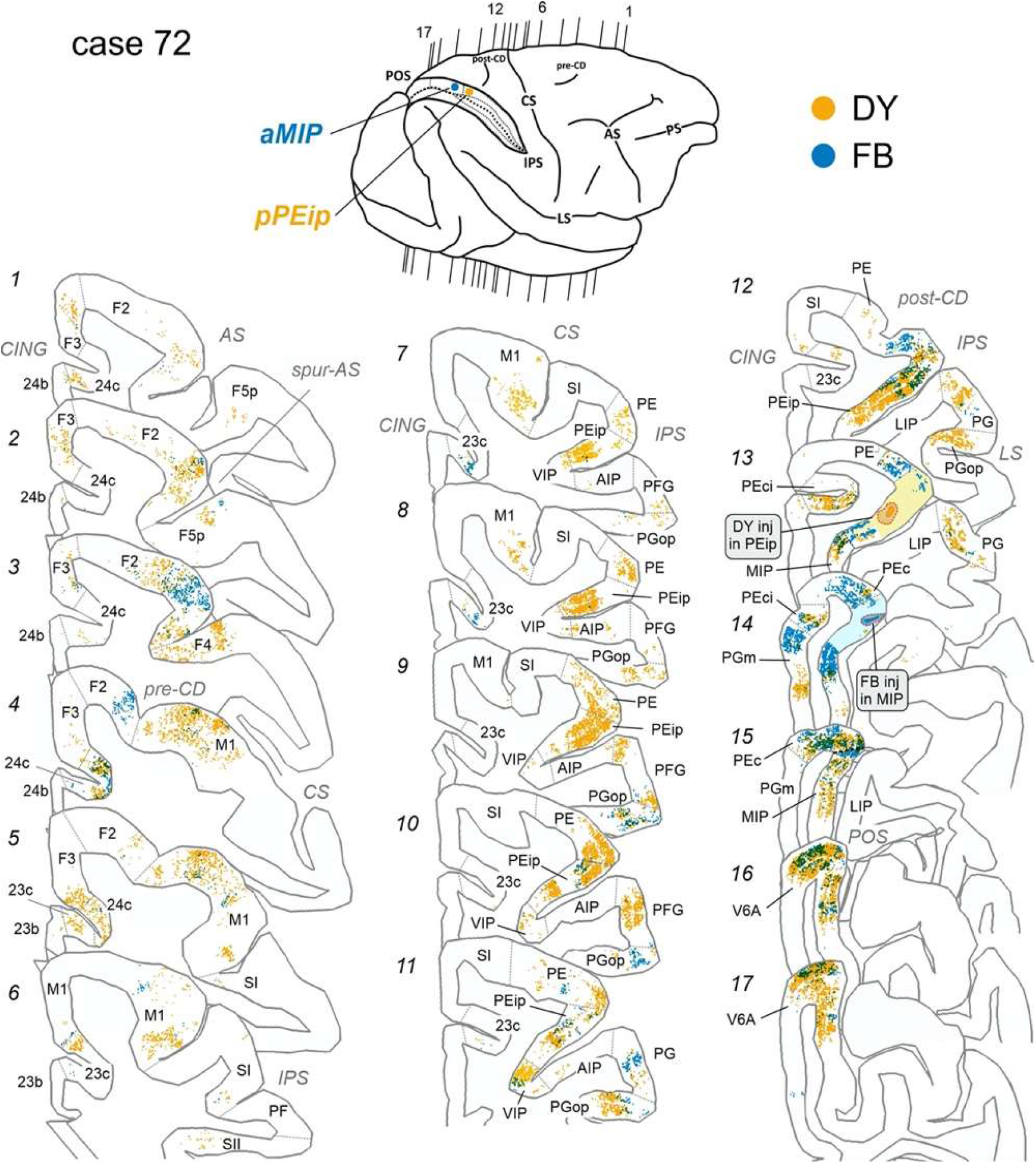
Distribution of retrogradely FB-labelled (shown in blue) and DY-labelled cells observed in Case 72 after the tracer injections in aMIP and pPEip, respectively, shown in representative sections through the frontal and the parietal cortex. The lightly colored zone surrounding the injection site in sections 13 and 14 corresponds to a sector with homogeneous intrinsic labeling. The levels at which the sections were taken is indicated in the drawing of the hemisphere in the upper part of the figure. POS = parieto-occipital sulcus; preCD and post-CD indicate pre-central and post-central dimple, respectively. Other abbreviations as in Figures 1 and 2.

**Fig. 5.**
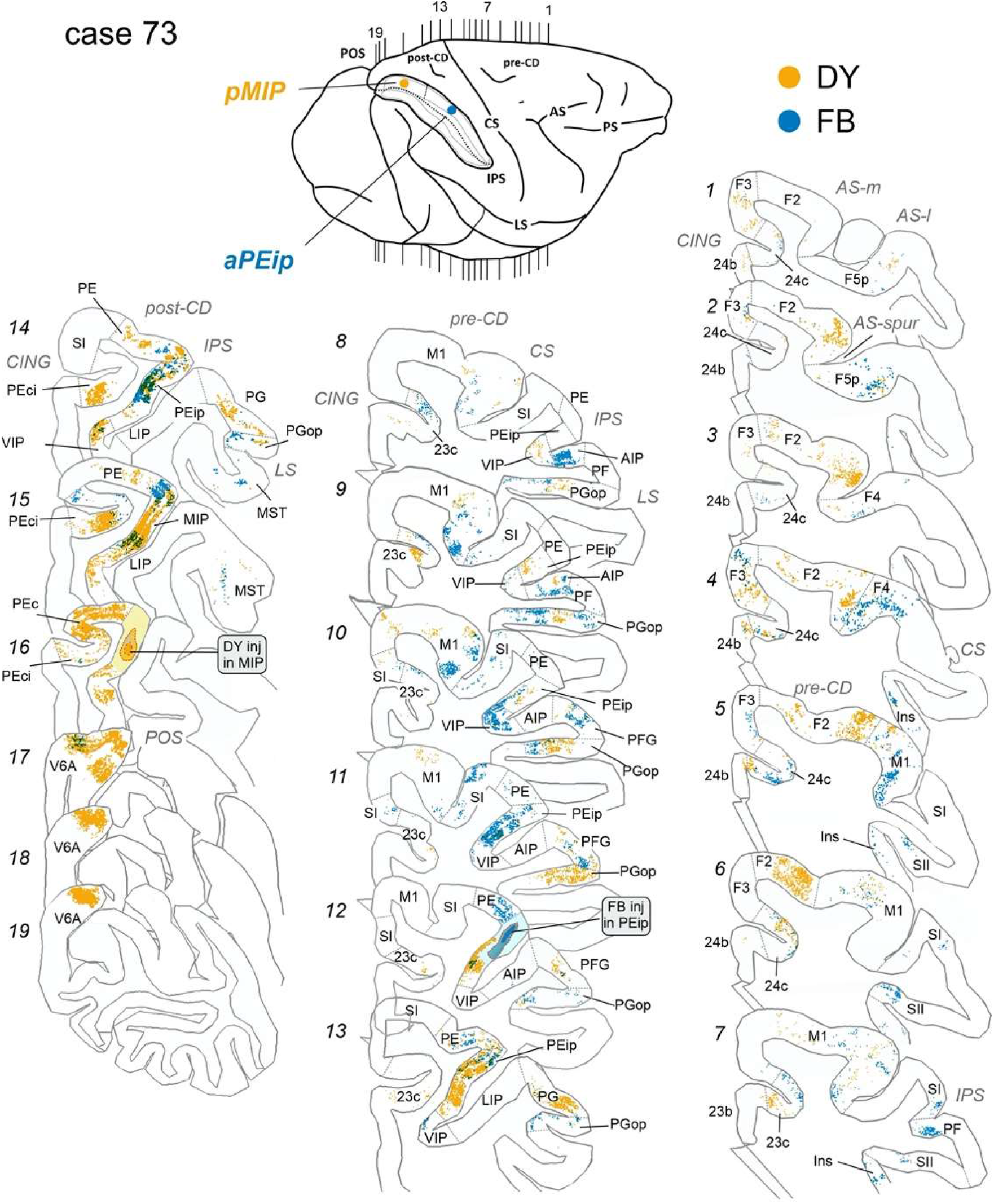
Distribution of retrogradely FB-labelled (shown in blue) and DY-labelled cells observed in Case 73 after the tracer injections in aPEip and pMIP, respectively, shown in relevant sections through the frontal and the parietal cortex. Conventions and abbreviations as in Figures 1,2 and 4.

##### Projections from frontal and cingulate cortex

In frontal cortex, RLC were found mostly in a region spanning from the ventro-rostral sector of area F2 (F2vr), around the spur of the arcuate sulcus, up to the border with M1 (primary motor cortex, F1) in the dorsal part of premotor cortex (Figs. 3, 4:*2-4*, 5:*2-6*). In both cases, they represented about 10% of the total number of RLC. Labelling extended over the classical arm region described in previous studies that combined anatomical tracing and physiological recording during reaching tasks (Caminiti et al., 1991; Johnson et al., 1996), as well as in the region of the arcuate spur, where neural activity is more related to hand movement (Fogassi et al., 1999). Smaller proportions of RLC (3,7-3,8%; Figs. 4: *4-6* and Fig. 5: *6-11*) were found over the arm region of M1 (see Johnson et al., 1996), lateral to the pre-central dimple. No RLC were found in the mesial part of M1, in the leg and foot representations, in line with data showing that neural activity in MIP is mostly related to visuomotor control of coordinated eye-hand actions.

A very small proportion of RLC was observed in area F3 (supplementary motor area, SMA; 1,3-1,6%; Fig 3), and a moderate number of them was located in the agranular cingulate area 24c/d (2-2,7% Figs. 4:*4-5*, 5:*4-6*) and in the granular cingulate area 23c (1,2-2,3%; Figs. 4:*7-8*, 5:7*-13*).

##### Projections from parietal cortex

In PPC, RLC were found in both the superior (SPL) and, to a lesser extent, inferior (IPL) parietal lobules. In SPL, after the aMIP injection, there was strong labelling in areas PEc (18,2%; Figs. 3,5: *14-15*), PEip (17,8%; Figs. 3, 5:*7-13*) and PE (13,8%; Figs. 3, 5: *10-12*), After the pMIP injection, the labelling was similarly robust in PEip (16,5%; Figs. 3, 5:*9-13*), weaker but still strong in PEc (12,5%; Figs. 3, 5: *3-16*), modest in PE (4,1%).

On the medial wall of the SPL, projections from area PEci were stronger to pMIP (12,9%) than to aMIP (6,1%; Figs. 3, 4:*13-14*, 5:*14-16*) and those from PGm were mostly addressed to aMIP (7,1%; Figs.3, 4: *14*). Finally, projections from area V6A were mostly (22.2%) addressed to pMIP (Fig.5:*17-19*), but in smaller proportion also to aMIP (7,3%: Fig. 4:*16-17*).

The only IPL areas projecting to MIP, although with a relatively modest proportion of cells (4,3% to pMIP; 3.65 to aMIP), were areas PG (Figs. 3, 5:*9-13*, 5:*13-14*) and PGop (Fig 4:*7-12*; Fig. 5:*8-13*). RLC were sparse in VIP (Fig.4:*7-11*), virtually absent in AIP, absent in LIP. Area MST contained a very small proportion (0,7%) of cells projecting to aMIP. Finally, very few. RLC were observed in SI and SII. No RLC projection to MIP were found in prefrontal areas.

#### Ipsilateral cortical projections to area PEip

Two tracer injections targeted PEip (Fig. 1), one in Case 73, where FB resulted to be placed at about its middle part, and one in Case 72, where DY was placed in the caudalmost part of it, adjoining the border with MIP (pPEip). As observed after the tracer injections in MIP, RLC substantially involved frontal and parietal areas, and their distribution reflected A-P gradients of connectivity in the db-IPS.

##### Projections from frontal and cingulate cortex

As shown in Table 1, after both the aPEip and the pPEip injections robust labelling was found in M1 (15,6% and 13,5%, respectively). Robust connectivity with M1, therefore, appears to be a unifying connectional feature of PEip, together with the projection to the spinal cord. In M1, the labelling was mostly located in the medial bank of the CS, thus involving the “new” M1 (Rathelot and Strick, 2009), where hand movements are represented (Figs. 4:*5-8*, 5:*5-10*). After the pPEip injection, RLC also extended more rostrally in M1 over the cortex of the precentral convexity, lateral to the pre-central dimple (pre-CD; Figs. 3, 4:*4-6*). Furthermore, after pPEip, but not aPEip injection, robust labelling was found in F2 (Figs. 3, 4:*1-5*). After the pPEip injection, the proportion of RLC in F2 (13,4%) was similar to that observed after that in aMIP (10,9%). However, RLC were almost completely located lateral to the pre-CD, whereas after the MIP injection they extended also more dorsally (Fig. 3). In both cases, moderate labelling also involved the ventral premotor area F4 (Figs. 3, *5:3*, 5:*4*) and weaker labelling was observed in F3 (Figs. 3, 4:*1-3*, 5:*4-5*). Moderate labelling was observed in areas 24c/d and 23c (Figs. 3, *4:1-7*, 5:*1-8*).

##### Projections from parietal cortex

In the SPL, robust labelling to both aPEip and pPEip was observed in area PE, richer after the aPEip injections (18,3% vs. 11,1%). In this area, RLC very densely packed in the rostral part, however after the pPEip injection they also extended in the caudal part, which was the PE sector densely labelled after the MIP injections (Figs. 3, 4:*7-12*, 5:*11-15*). Caudal to PE, after the pPEip injections, labelling was relatively moderate in PEc (4,2%) and PEci (5,3%), weak in PGm (1,6%), and robust in V6A (10,5%; Figs. 3, 4:*13-17*). In all these areas, labelling was much weaker, or even absent after the aPEip injection (Figs. 3, 5:*14-19*). Similarly, the number of RLC observed in MIP was much higher after the pPEip (12,9%) than the aPEip (5,1%) injection. In the IPL, both aPEip and pPEip were moderately connected with the hand-related area PFG, though after the pPEip injection the labelling moderately involved also PG (Figs. 3, 4:*7-13*, 5:*5-7*). Furthermore, aPEip was characterized by a robust input from PGop (11,7%; Fig.4:*8-10*), which was much weaker for pPEip (4,2%), as well as by relatively robust input from the hand-related area AIP (6,3%) and in VIP (5,5%), where RLC were relatively sparse after the pPEip injections (Figs. 3, 4:*8-12*, 5:*8-14*).

After the aPEip injection there was robust labelling in SI (7,3%; Figs. 3, 5: *6-7*) and a relatively weak labelling in SII and the insular cortex, all virtually devoid of labelling after the pPEip injection. Finally, a weak input from MST was observed in both cases.

#### Connectivity profiles of aPEip, pPEip, aMIP, and pMIP

To offer a quantitative view of the results, the data reported in Figs. 4-6 and in Table 1 were expressed in the form of frequency distribution. Figure 7 reports the proportion of RLC (Y axis) across cortical areas, which are arranged from left to right (X axis) according to their approximate A-P location in the cortex.

**Figure 6.**
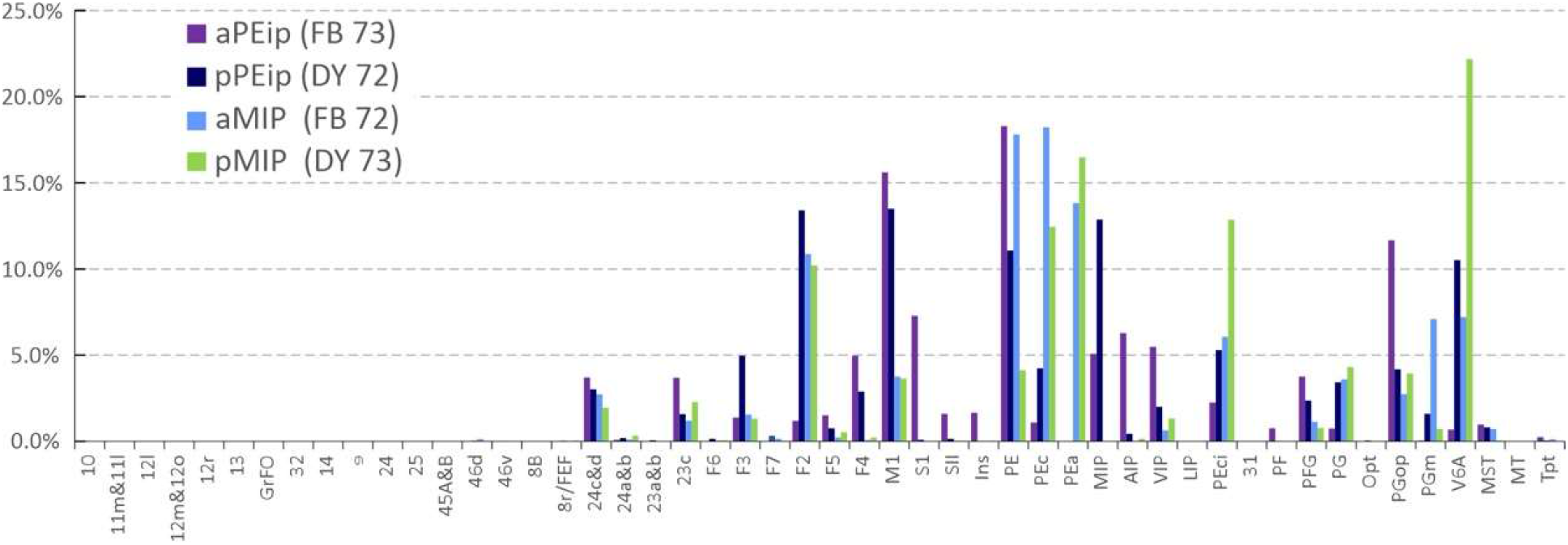
Ipsilateral cortical projections to areas aPEip, pPEip, aMIP, pMIP. Proportion of cells projecting from different areas to the four injection sites located in area aPEip (violet), pPEip (dark blue), aMIP (light blue), pMIP (green). pMIP cells projecting to PEip, and vice versa, are included. Percentages are calculated relative to the total counts of RLC obtained after each injection.

**Figure 7.**
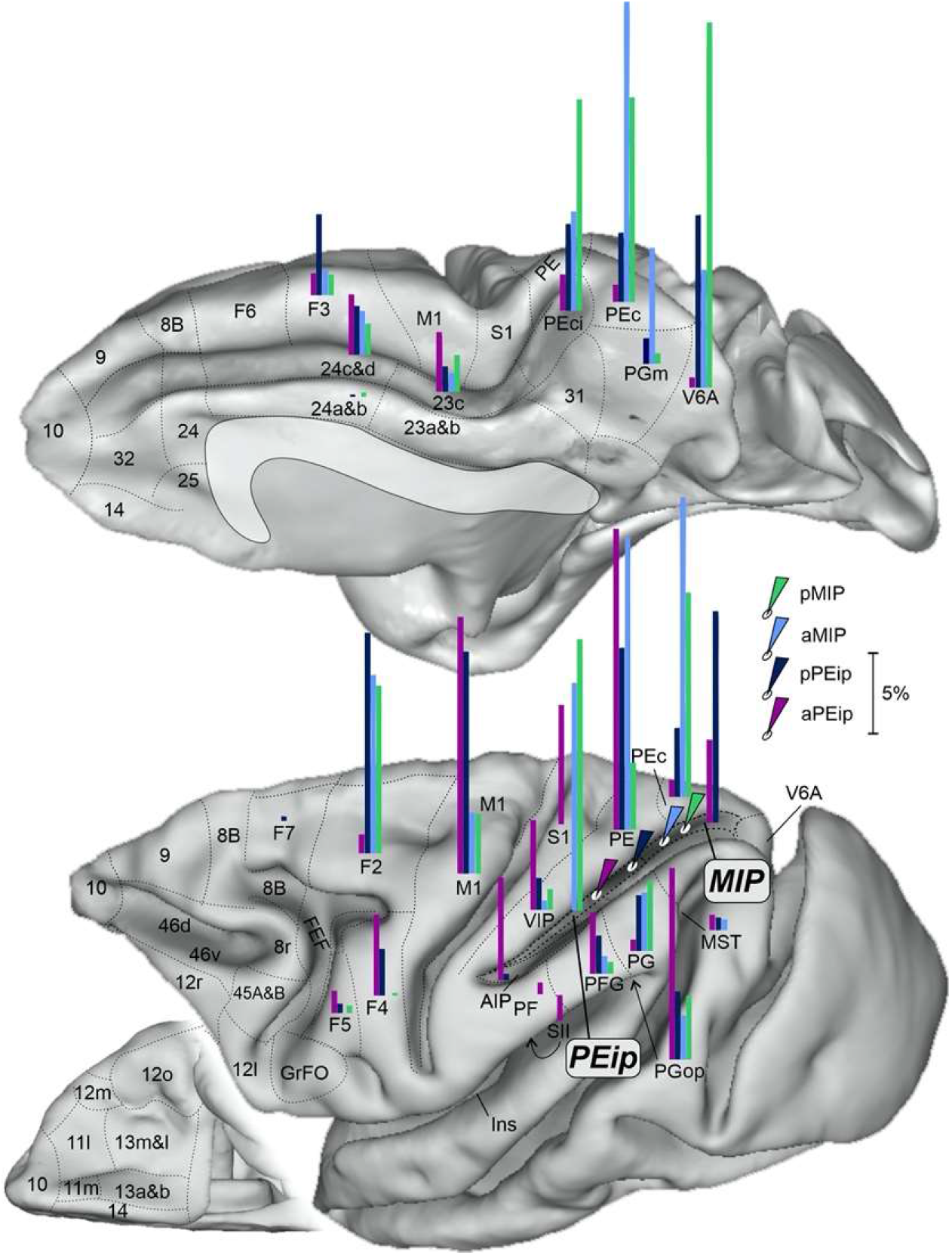
Gradient-like organization of the parietal and frontal projections to the dorsal bank of the IPS. Mesial (top), lateral (bottom, right) and ventral (bottom, left) views of the monkey brain showing the proportion of projecting cells (see Fig. 6) in their relative anatomical location, after tracer injections (white ovals with colored arrows) at the four A-P levels of the db-IPS. Each bar has a length proportional to the percent of RLC (range 1-30%, scale bar corresponding to 5%) to aPEa (purple), pPEa (dark blue), aMIP (light blue) and pMIP (green). Conventions as in previous figures.

The frontal input to parietal areas injected in this study stems mostly from areas F2 and M1. Projections from F2 are mainly addressed to pPEip, aMIP and pMIP, in decreasing order of magnitude. Motor cortex projections follow a similar pattern but differ for a strong input to aPEip as well. Area S1 projects only to aPEip. Smaller projections stem from cingulate areas 24c and 23 and from ventral premotor area F4.

The parietal projections to PEip and MIP are by far stronger that the frontal ones and originate mainly from superior parietal areas, such as PE, PEc, from local connections within PEip and MIP and from V6A, PEci, and PGm. Inferior parietal projections are by far weaker, and originate from PGop, especially after the injection in aPEip, with smaller contribution from areas PG and PFG. Finally, aPEip showed a relatively robust connection with areas AIP and VIP.

In several instances, the projections addressed to areas PEip and MIP from cingulate, frontal and parietal areas followed a gradient-like pattern, as also shown in Fig. 8. Examples are the projections from area 24c, M1, and PFG, which all project with decreasing strength to aPEip, pPEip, aMIP and pMIP. The F2 projections to dorsal intraparietal areas display a similar pattern, if one excludes the scant projection to aPEip. On the contrary, the strength of PEci projections shows an inverse gradient. The strength of the projections from PE and V6A waxes and wanes in the A-P extent.

**Figure 8.**
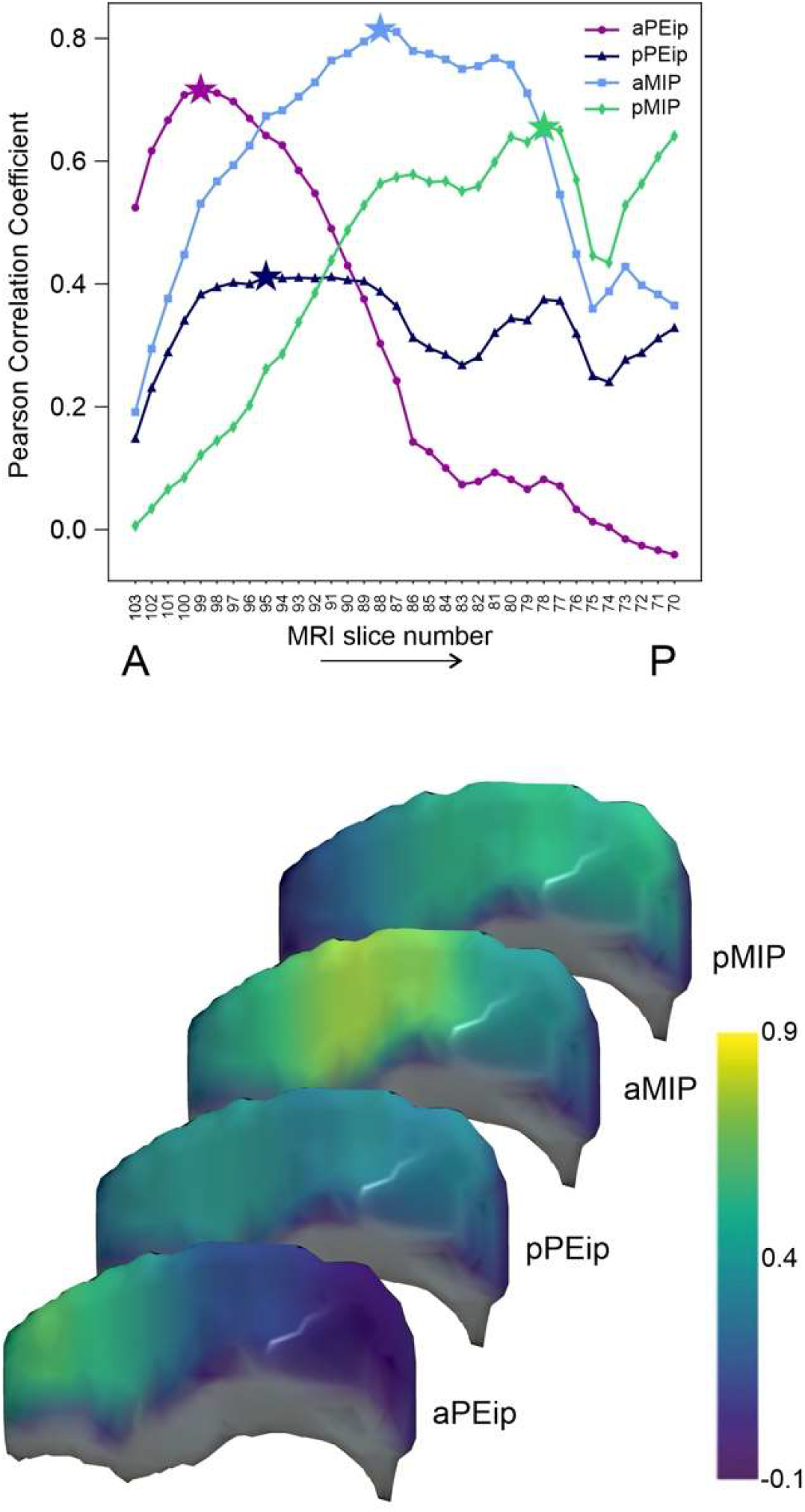
**A**. Pearson’s correlation coefficient between the distribution of diffusion-based connectivity estimated in 2.5 mm windows along the db-IPS and the distribution of labelled cells after the four injection in aPEip, pPEip, aMIP, pMIP. MRI slice numbers refer to the central position of each sliding window, where slice 103 is anteriormost and slice 70 the posteriormost. The star markers indicate the A-P location with the highest correlation coefficients. **B**. The Pearson’s correlation coefficients after each of the four injections are also reported in colour code across the db-IPS. Colour bar on the left.

A pictorial representation of the gradient-like organization of this part of the parieto-frontal system can be seen in the brain figurine of Figure 6.

#### Segregation and overlap and laminar distribution of frontal and parietal cells projecting to PEip and MIP

In the tangential domain of the cortex there exists an orderly arrangement of properties that can relate to the representation of sensory receptors, motor output, visual attention, motor intention, working memory, etc. Moreover, there is evidence that cortical connections shape, at least in part, the functional properties of neurons in the parieto-frontal system (Johnson et al., 1996; Chafee and Goldman-Rakic, 1998, 2000; Battaglia-Mayer et al., 2001).

To study whether PEip and MIP share cortical afferents, therefore functional properties, we compared the tangential distribution of frontal and parietal cells projecting to their anterior and posterior sectors, a study made possible by the injections of two different fluorescent tracers in each of the two animals.

In case 72, where DY was injected in pPEip and FB in aMIP, frontal cells projecting mostly to pPEip (Fig. 4, see yellow labelling) involve both dorsal premotor area F2 and M1 while those projecting to aMIP (Fig. 4, see blue labelling) occupy restricted efferent frontal zones, mainly located in F2. With the exclusion of a restricted part of the latter (Fig. 4:*2-3*), cell projecting to pPEip and aMIP were largely segregated in the tangential domain of the cortex. At some locations, parietal cells projecting to both pPEip and aMIP were segregated (Fig.4:*7-17*), even in the same area, as for PGm (Fig.2:*14*). On the contrary, extensive overlap was found in areas PEc, PEci and V6A (Fig. 4:*14-17*).

The distribution of cells projecting to aPEip and pMIP, where FB and DY were respectively injected (Fig. 5) obeys to a similar pattern, where segregation dominates over overlap in both frontal and parietal projections, although some overlap was observed in areas PGop (Fig.5:*10-11*), pPEip (Fig. 5:*13-15*), aMIP(Fig. 5:*15*), V6a (Fig. 5:*17*).

When comparing the distribution of cells in the rostral bank of the CS, i.e., in the “new M1” (Rathelot and Strick, 2009), in both cases 72 and 73 we mostly observed absence of overlap of cells projecting to the intraparietal areas injected, as well as in area PE and in large part of aPEip, while a small overlap was confined only to very limited zones of the bank (Fig. 5:*6-7*). Finally, the analysis of the laminar distribution of RLC in the various frontal and parietal areas more densely labeled after the injections in different sectors of PEip and MIP showed a proportion of RLC in the superficial vs. deep layers virtually everywhere within 33% and 66%, that is a marked bilaminar distribution.

### DW-MRI study and comparison with histological tracing

To compare the IPS connectivity obtained through histological procedures, as reported above, with that obtained through DW-MRI, we computed the Pearson’s correlation coefficient between the distribution of RLC obtained after each injection sites and the distribution of diffusion-based connectivity obtained from the intra-axonal MRI signal fraction estimation (see Material and Methods) at different locations along the entire extent of the db-IPS. In other words, we compared the proportion of RLC cells of any of the 48 cortical regions in which they were found (see Material and Methods) with the diffusion-based connectivity shown by these areas to different IPS MRI sections (38 in totals, see Methods). To cover in a continuous fashion the whole IPS we have used a sliding window of 2.5mm, corresponding to five MRI coronal slices, moving from anterior to posterior (A-P) and selecting all streamlines connecting the MRI slices to the 48 cortical ROIs included in our analysis (see Material and Methods). To better reproduce the extent of the injection sites of retrograde tracers in the dorso-ventral dimension, the MRI slices encompassed only the dorsal and middle sectors of our three-fold subdivision of the db-IPS (Fig. 8B).

In Figure 8A data points in each curve show the Pearson’s coefficients for the correlation between the distribution of RLC obtained for each of the injection sites (aPEip, pPEip, aMIP, pMIP) and the diffusion-based connectivity of each 2.5mm sliding window along the A-P dimension of the db-IPS. The X-axis shows the MRI coronal slice number at the center of each window. The locations with the highest correlation are indicated by the star markers. The MRI coronal slice number corresponding to each injection site’s highest correlation coefficient (Fig. 8A; star markers) well agrees with the relative position of the injection sites of neural tracer (Fig. 1). In fact, the RLC distribution after injection in aPEip had the highest correlation value (r=0.72; n=34; p=1.1*10^−8^) at slice 99, after injection in pPEip at slice 95 (r=041; n=48; p=0.004), showing however similar correlation values (plateau) at different A-P locations ranging from slice 97 to 89, while after injection in aMIP the correlation peaked at slice 88 (r=0.81; n=34; p=1.9*10^−12^) and after injection in pMIP at slice 78 (r=0.66; n=34; p=3.9*10^−7^).

When selecting the locations with highest correlation for each of the four injection sites, the overall correlation between the diffusion-based connectivity estimation and the RLC distribution was r=0.65 (n=192, p=1.7*10^−24^).

The changes of the correlation coefficient between the distributions of labelled cells and diffusion connectivity across the db-IPS are shown in Figure 8B. It can be seen that the highest correlation was found in a region spanning the central (in A-P dimension) and dorso/middle (in dorsoventral dimension) sectors of the bank, after injections in aMIP. A good correlation was also found in the anterior third of the bank after injections in aPEip, while the correlation decreased, although to a different extent, after injections in pPEip and pMIP.

The corresponding distribution of RLC for the four injection sites alongside the diffusion-based connectivity for the locations with the maximum Pearson’s coefficients are reported in Figure 9, together with the relative MRI slices and drawing of the histological sections.

**Figure 9.**
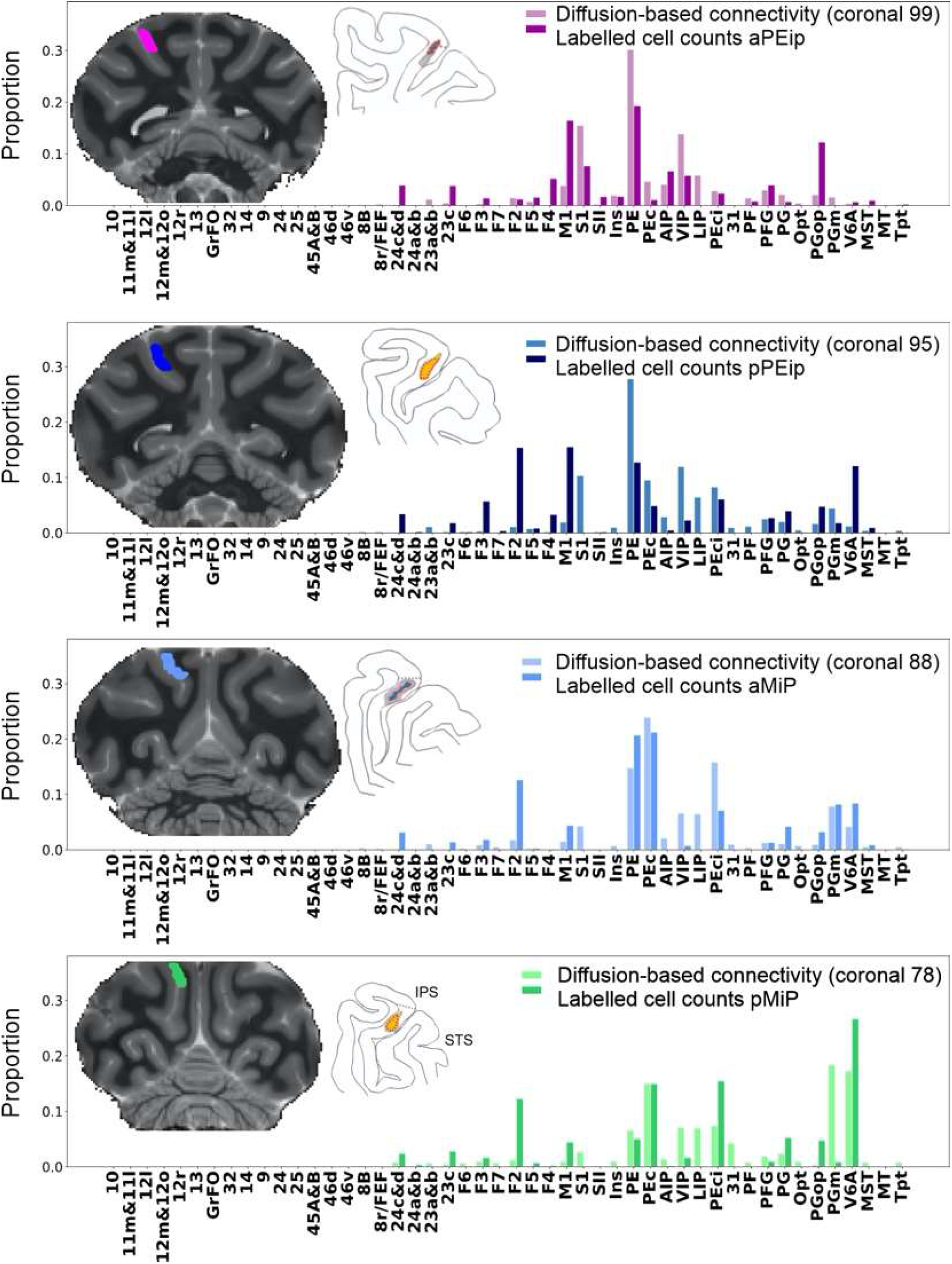
Distribution of labelled cells and diffusion-based connectivity for locations with maximum Pearson’s correlation coefficients (aPEip: r=0.72; pPEip: r=0.41; aMIP: r=081; pMIP: r=0.66). For each distribution, the MRI slices corresponding to the center positions of the sliding windows with highest Pearson’s correlation coefficients are reported next to the reconstruction of the histological sections where the injection sites were found. The local connections between MIP and PEip are not reported.

For the four injections sites there are 192 (48 areas x 4 injections) potential ROIs connections, among which 113 have non-zero labelled cell counts. Diffusion tractography shows an average of 90.4% of the connection’s weights for ROIs with non-zero reported labelled cells. Moreover, tractography correctly identified 108 connections (true-positive connections; TP), thus missing only 5 connections (false-negative connection; FN). Tractography correctly reported no connectivity for 44 ROIs (true negative connections; TN), but estimated connectivity for 36 ROIs where no labelled cells were found (false positive connections; FP). Overall, this resulted in a sensitivity of 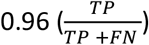 and a specificity of 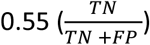.

Moreover, we evaluated in a quantitative fashion the degree of similarity of the diffusion-based connectivity estimation along the db-IPS. To this aim, we computed the Pearson’s correlation coefficient between all sliding windows.

Figure 10A shows the Pearson’s correlation coefficient between the distributions of diffusion-based connectivity estimated in different sliding windows along the A-P extent of the db-IPS. The X and Y axes show the MRI coronal slice number corresponding to the center of each window. A strong correlation is expected between locations distant four or less MRI slices apart, due to the windows overlap. A decrease in correlation can be observed when the distance between windows increases in the A-P extent of the bank. This suggests a general gradient-like organization, where the pattern of cortical connectivity gradually changes. Visual inspection of the correlation matrix highlights the existence of three potential clusters, located anteriorly, centrally and posteriorly along the bank, that can be identified by their highest correlations (range 1-0.6) between neighboring locations. This suggests that along the A-P extent of the db-IPS there might exist three broad connectionally different regions. A similar matrix (Fig. 10B) is shown for selected locations corresponding to the four MRI windows with the highest correlation between the diffusion-based and tract tracing connectivity (see also Fig. 8). It can be seen that similar results were obtained when correlating the pattern of connectivity obtained from histological tracing data, after injections in intraparietal areas aPEip, pPEip, aMIP, and pMIP.

**Figure 10.**
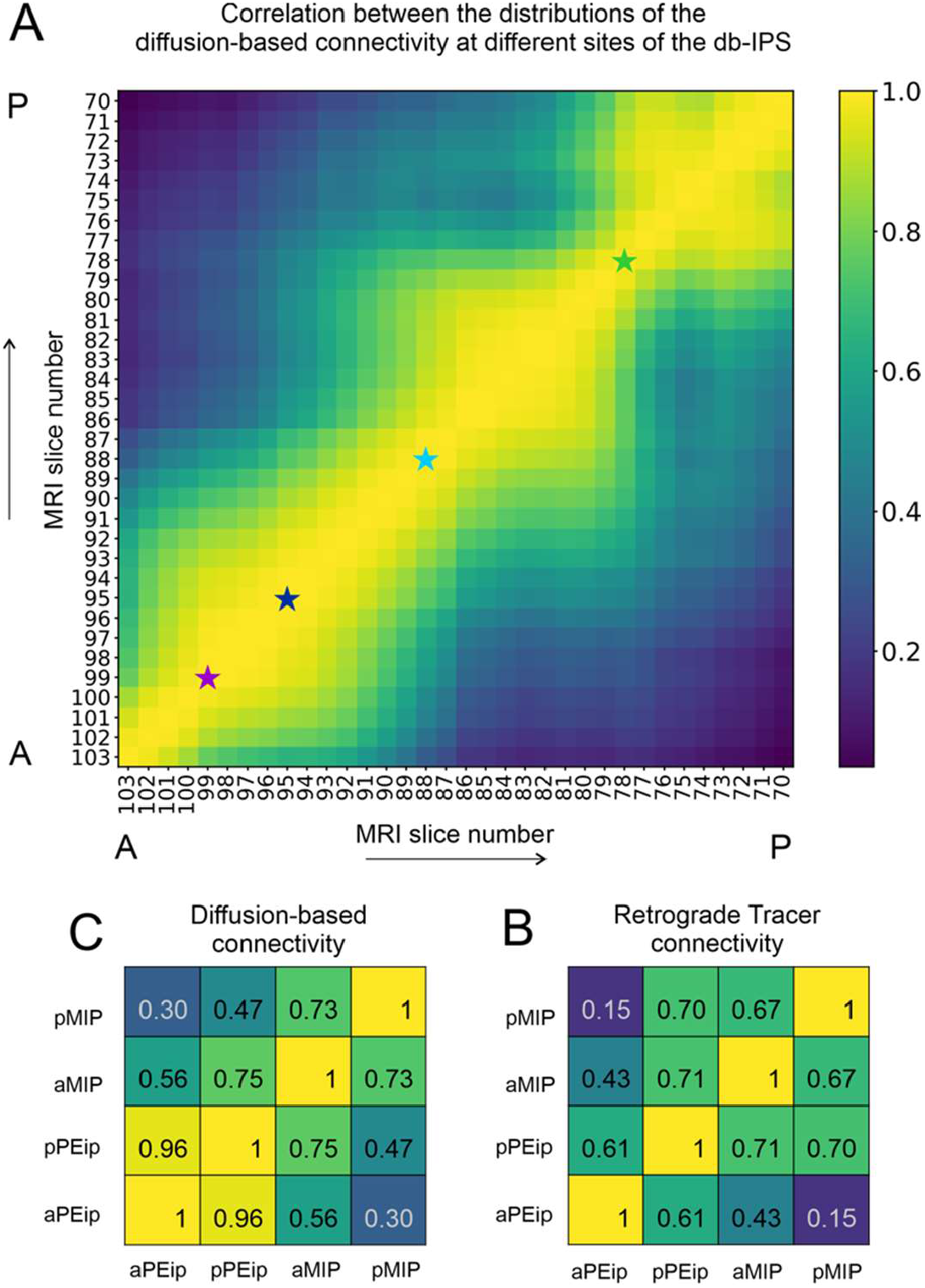
**A**. Pearson’s correlation coefficient between the distributions of the diffusion-based connectivity estimated in subregions along the db-IPS, as defined by a sliding window of 2.5mm moving in the anterior-posterior direction (5 MRI coronal slices). For each window, the connectivity is evaluated first by selecting all the streamlines connecting the MRI slices to the 48 ROIs included in the analysis and summing the contribution to the intra-axonal MRI signal fraction of each streamline for each cortical area. Data were normalized relative to the total contribution of the streamlines associated to each sliding window. The X and Y axes show the MRI slice number corresponding to centre position of each window. Star markers (slices 99, 95, 88 and 78) indicate the locations with highest correlation coefficient between diffusion-based connectivity and labelled cells, after tracer injections in aPEip, pPEip, aMIP, and pMIP (see Fig. 8). Values of correlation coefficients are indicated by the colour code (see bar on the right). **B**. Pearson’s correlation coefficients between the distributions of diffusion-based connectivity estimated at the four sites reported above. **C**. Pearson’s correlation coefficients between the distributions of RLC after injection in aPEip, pPEip, aMIP, pMIP. In **B** and **C** correlation coefficients are also reported with relative values (colour code as in **A**).

Furthermore, we investigated the cortical connectivity of the dorsal, middle and ventral sectors of the db-IPS using diffusion MRI. It is worth stressing, the cortical regions lying in the more ventral and deep part of the bank can be hardly accessed by neural tracer injections, therefore their connectivity remains virtually unknown. The sum of the diffusion-based connectivity calculated across the 38 different A-P locations (MRI slices) for the dorsal, middle and ventral sectors is shown in Figure 11. The parietal areas VIP, V6A, PE, LIP, PEc, PGm, and SI are the ROIs showing the overall strongest connectivity with the bank, among the 48 ROIs considered in this study.

**Figure 11.**
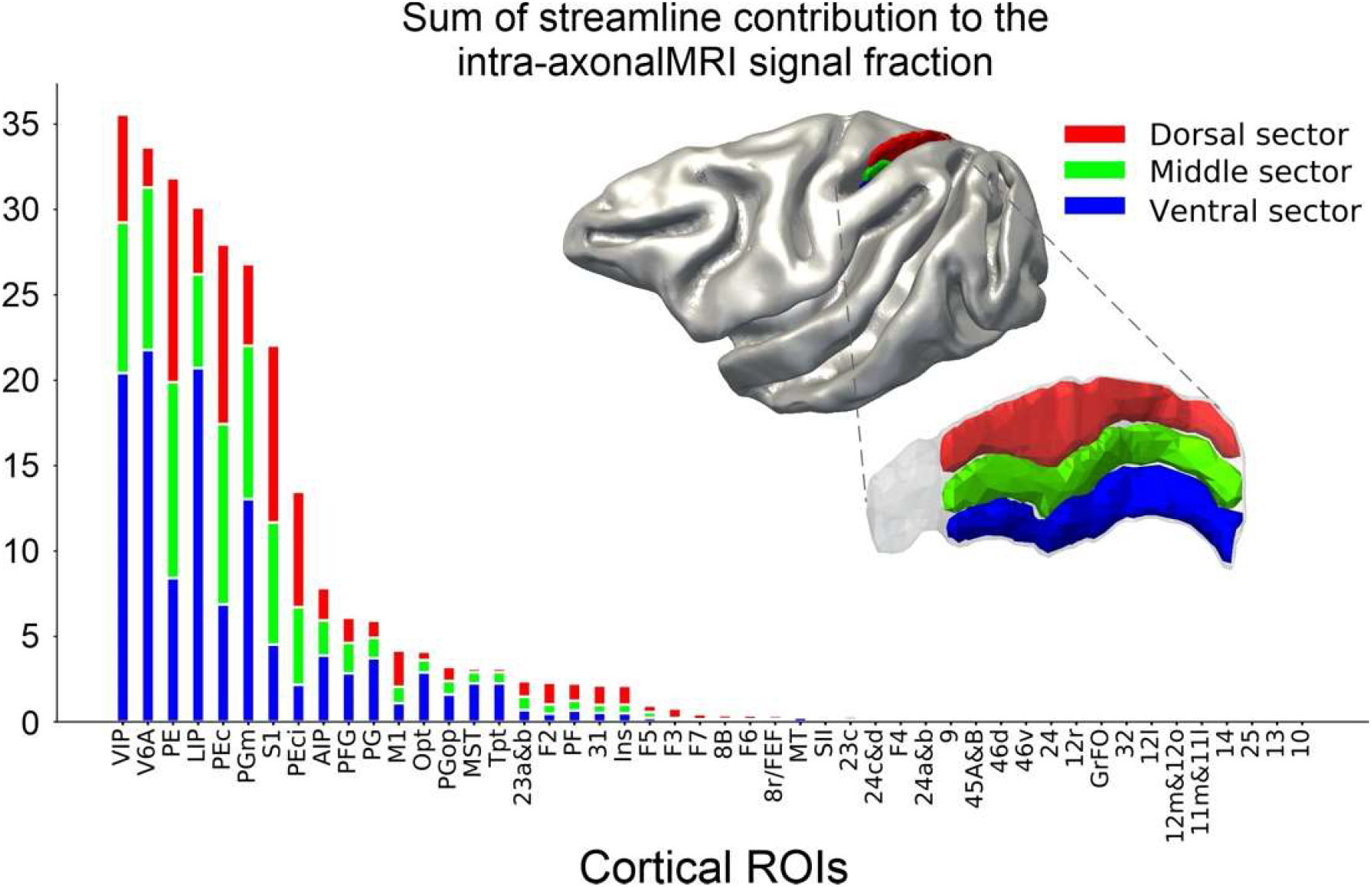
Sum of the cortical connectivity of the db-IPS to other cortical ROIs. For each ROI, the diffusion-based connectivity estimation is reported for the dorsal (red), middle (green) and ventral (blue) sectors. The diffusion connectivity corresponds to the sum of streamline contributions to the intra-axonal MRI signal fraction estimated using COMMIT for each cortical ROI. The sectors of the db-IPS are shown on the mid cortical surface (top right) and on the db-IPS (bottom right).

However, clear differences emerge in the streamline contribution provided by specific portions of the IPS along the dorso-ventral dimension. To highlight this aspect, we report the results (Fig. 12) referring to the connectivity occurring between each of the 12 most connected cortical areas (i.e., VIP, V6A, PE, LIP, PEc, PGm, SI, PEci, AIP, PFG, PG, M1; see Fig. 11), and the A-P and D-V extent of the db-IPS. Each image shows the spatial distribution of the diffusion-based connectivity, along the 38 A-P dorsal, middle and ventral subdivisions of the bank, for each of the 12 cortical ROIs listed above. The sectors displaying strong connectivity with the indicated cortical ROI are shown in yellow and orange. It can be seen that there exists a smooth transition in the strength of connectivity in both the A-P and D-V dimensions of the bank. The IPS region more strongly connected with area VIP is the most anterior sector of the bank, with a gradual reduction moving posteriorly, while for V6A is the postero-ventral part of the bank, as also observed from tract tracing data on the proportion of RLC (see Fig. 7). Area PE instead display a more diffuse pattern of connectivity along the D-V dimension of the anterior part of the bank. LIP connectivity occurs exclusively with the regions located in the more ventral part of the dorsal bank, close to the fundus of the IPS. Another example of a gradient-like distribution of connectivity, along both the A-P and D-V dimensions is offered by PEc, whose connectivity is strongest with the dorsal and intermediate part of the bank. The connectivity of PGm resembles that of V6A, but it is weaker and more diffuse in the A-P extent of the ventral part of the intermediate sectors. Area SI is strongly connected with the D-V extent of the rostralmost part of the bank, while the connections of PEci are more selective, since they occur mainly with the central part of the bank, are stronger dorsally and fade away moving ventrally, anteriorly and posteriorly. (see Table 2). The inferior parietal areas AIP, PFG, and PG show a weak connectivity with the anterior part of the ventral sector of the bank, while motor cortex (M1) is weakly connected with its antero-dorsal sector.

**Figure 12.**
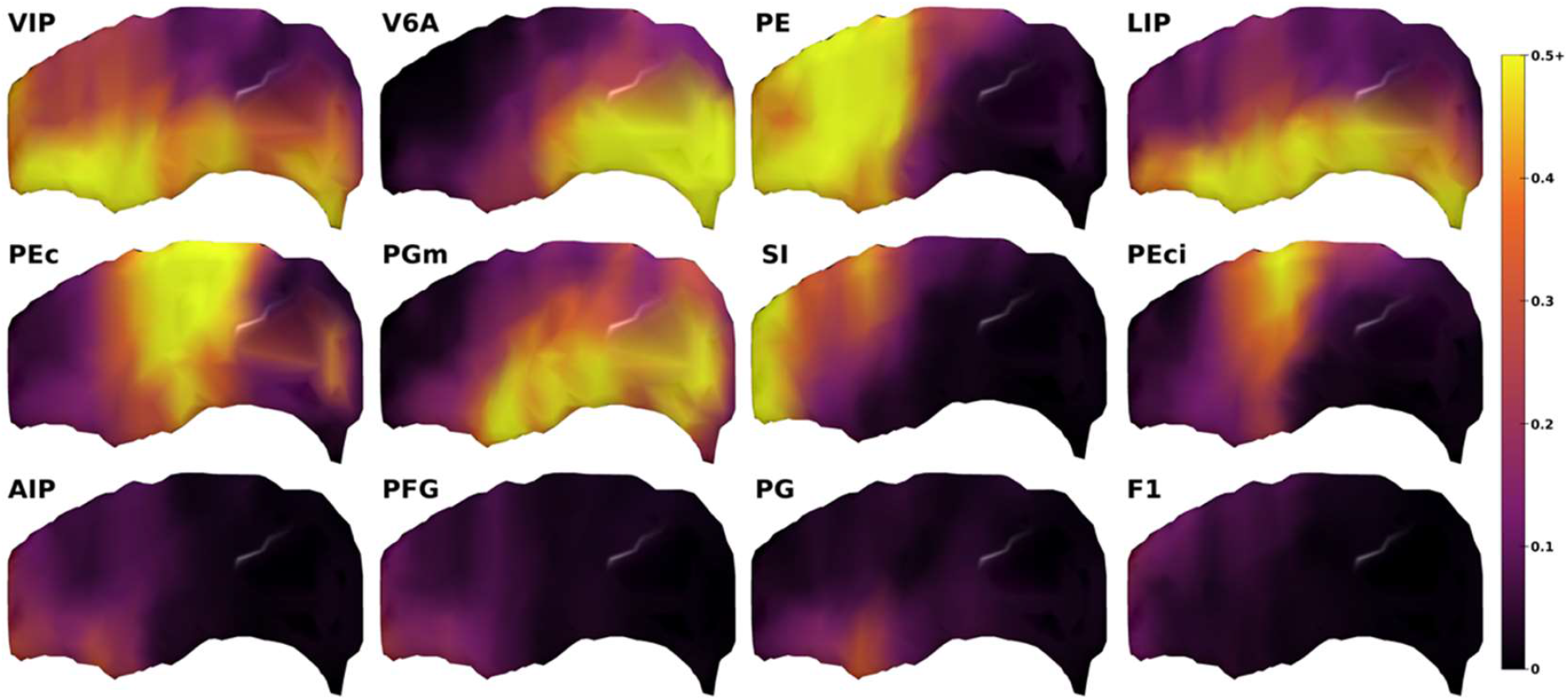
Spatial distribution of the IPS connectivity estimated from DW-MRI along 34 dorsal, middle and ventral anterior-posterior sectors of the db-IPS, for the 12 cortical ROIs displaying the strongest estimated connectivity with the db_IPS (see Fig. 11).

## DISCUSSION

This study combined histological tract tracing and diffusion tractography data to highlight the connectivity of the db-IPS, including its the deepest part where injecting selectively neural tracers remains difficult.

Altogether, the present data provide solid support for the subdivision of the db-IPS into a rostral area PEip and a caudal area MIP, based on the distribution of corticospinal neurons, as well as for an internal subdivision of both areas into an anterior and posterior sector. However, the data also show antero-posterior and dorso-ventral connectional gradients, matching the gradients of functional properties described by electrophysiological studies. Ealier studies had reported differences in the functional properties of cells in the A-P extent of the SPL (Crammond and Kalaska, 1989; Burbaud et al., 1991), suggesting that activity in area PE is more related to somatosensory function, while in MIP (Colby and Duhamel, 1991) it is more related to motor and visual functions. A combined anatomo-functional analysis of this region (Johnson et al., 1996) in behaving monkeys revealed that reaching-related neurons displaying signal-, set-, movement- and positional-related activity were encountered more frequently moving from the ventral (therefore posterior) part of MIP to its dorsal part and to area PE. A mirror-image trend was found in the frontal lobe, where the proportion of arm position- and movement-related neurons decreased and that of set- and signal-related neurons increased moving from motor cortex posteriorly to dorsal premotor areas (F2/F7) anteriorly. This same study revealed that parietal and frontal regions displaying similar activity types were linked by direct cortico-cortical connections, suggesting the latter contribute in shaping the dynamic properties of cortical neurons across distant areas.

### Cortical connections of the db-IPS

The present data are in line with, but also extend data from the only previous study (Bakola et al., 2017) focused on the connectivity of MIP vs. the parieto-occipital area V6A. Noteworthy, in that study, MIP defined myeloarchitectonically extends rostrally up to the A-P level of the caudal end of the central sulcus, thus including the caudal part of the corticospinal sector of the db-IPS.

Indeed, our data from tracer injections in MIP show a relatively strong connectivity with visuomotor areas V6A and PEc with PEip, and both the peri-precentral dimple and ventrorostral parts of area F2. Weaker connections involve the IPL visuomotor area PG, area PGop and M1. Furthermore, aMIP, when compared to pMIP, shows a stronger connectivity with the somatosensory area PE and visuomotor area PGm and a weaker one with somatosensory area PEci. This connectivity pattern of MIP is substantially in line with that reported by Bakola et al (2017) for the caudal part of this area. Furthermore, indirect support for this connectivity pattern and for the reciprocity characterizing connections of MIP comes from studies in which this area was labelled after retrograde tracer injections in V6A (Marconi et al., 2001; Gamberini et al., 2009; Passarelli et al., 2011), PEc and PE (Marconi et al., 2001; Bakola et al., 2010; 2013), PGm (Passarelli et al., 2018), PG (Rozzi et al., 2006) and F2 (e.g., Johnson et al., 1996; Matelli et al., 1998; Marconi et al., 2001; Tanné et al., 2002). Thus, the connectivity pattern of MIP provides a neural substrate for the visuomotor control of reaching and eye-hand coordination, since it can serve as interface between the premotor areas of the frontal lobe and the parieto-occipital areas V6A and PEc, where neurons combine in a directionally-congruent fashion eye- and hand-related positional- and movement-related signals within their directional tuning fields (Battaglia-Mayer et al., 2000, 2001). Interestingly, similar inputs to MIP come from PGm (7m), where individual neurons also combine visual, eye and hand related signals (Ferraina et al., 1997a, b).

A model relevant to eye-hand coordination (Mascaro et al., 1983) integrating inputs from the retinal position of the target with eye- and hand position shows that both feedforward and recurrent interactions of these signals account very well for the experimentally observed tuning fields of parietal neurons. In this model, surprisingly the representation of directional variables concerning hand and eye movement emerges from Hebbian synaptic plasticity alone (for an overview on the network subserving eye-hand coordination see Battaglia-Mayer and Caminiti 2002; Battaglia-Mayer et al., 2015; Caminiti et al., 2017; Battaglia-Mayer and Caminiti, 2017).

Our data also show that area PEip as defined in the present study is a db-IPS sector displaying as unifying connectional features robust connectivity with the cervical spinal cord and the hand field of M1. Strong connections with area PE and with visuomotor hand-related area PFG (Ferrari-Toniolo et al., 2015), bimodal visual and somatosensory area VIP, and area F4 further characterize PEip. The caudal part of PEip also displays connections with V6A and F2 and a connectivity pattern with areas PEci, PEc, and PG quantitatively more similar to that of aMIP. In contrast, aPEip displays connections with the arm/hand field of SI, the hand-related area AIP and a strong connectivity with PGop. The connectivity pattern observed after the tracer injections in pPEip and aPEip is very similar to that observed by Bakola et al. (2017) after an injection in rostral myeloarchitectonic area MIP and in area PEip, respectively. Connections with PEip have been observed after retrograde tracer injections in areas V6A (Gamberini et al., 2009), PE (Bakola et al., 2013), PFG (Rozzi et al., 2006), AIP (Borra et al., 2008; Lanzilotto et al, 2019), F2 (e.g., Johnson et al., 1996; Matelli et al., 1998; Tanné et al., 2002) and M1 (Strick and Kim, 1978; Matelli et al., 1986; Hatanaka et al., 2001). This connectivity pattern fits very well with a possible role of PEip in sensorimotor control of hand movements. Indeed, area PEip as a whole appears to coincide with the db-IPS sector hosting corticospinal neurons making di-synaptic contacts with distal hand muscles motorneurons (Rathelot et al. 2017). Furthermore, this same sector seems to correspond to that part of the db-IPS hosting neurons with somatosensory receptive fields on the hand (Iwamura et al., 1994; Iwamura 2000; Seelke et al., 2012). The posterior part of PEip could also correspond to the sector hosting neurons with bimodal, visual and somatosensory receptive field centred on the hand (Iriki et al., 1996) and the anterior part to the sector where Gardner et al. (2007) recorded grasping-related neurons. The connectional differences between the posterior and the anterior part of PEip, suggest for the former a role in visuo- and somato-motor control of hand and, possibly arm movements, and for the latter a role in somato-motor control of hand actions.

### Diffusion-based connectivity estimations

In this work, we have used state-of-the-art DW-MRI Particle Filtering Tractography (PFT) algorithm with the COMMIT microstructure estimation method to examine the intra-axonal MRI signal fraction associated with streamlines, as opposed to directly use the number of streamlines. Such approach allowed to reduce density biases associated with white matter bundle features, such as length, curvature, and size, and make tractography more quantitative (Daducci, et al. 2014; Girard et al., 2014). This was achieved by using a model of the tissue microstructure (Stick-Zeppelin-Ball model, Panagiotaki et al., 2012, Daducci et al., 2014) ideal to explain the measured DW-MRI signal from the streamlines, by removing or penalizing redundant or inaccurate trajectories. In a previous study, Girard et al. (2020) compared various diffusion-based connectivity estimation approaches in the monkey brain and showed that *PFT-COMMIT* had strong performances in the prediction of parieto-frontal binary connectivity (sensitivity and specificity). Moreover, it had the highest fraction of valid connectivity weight among methods with high sensitivity and specificity.

In the connectivity network emerging after the four injection made within the dorsal bank of the intraparietal sulcus, our tractography results showed an increased sensitivity of 0.96 (from 0.79) and a decreased specificity of 0.55 (from 0.60), as compared to the more extensive analysis of the parieto-frontal network we made before (Girard et al., 2014). Overall, this resulted in an increased Youden’s index (Sensitivity + Specificity – 1; Youden, 1950) to 0.51 vs. the 0.39 reported in Girard et al. (2020). Moreover, in the network studied here, we found 90.4% of the connectivity weights between ROIs with reported non-zero labelled cell count, 10.2% more than reported by Girard et al. (2020). This suggests a strong predictive power of tractography for the connectivity of the monkeys IPS, which was also confirmed by the lack of connections with prefrontal areas shown by both histological and tractography results.

Furthermore, the connectivity estimated by tractography showed very clear antero-posterior and dorso-ventral gradients along the dorsal bank of the IPS, as also shown by histological studies, confirming that the gradient-like organization emerges regardless of the methodological approach used and might be a general principle of cortico-cortical connectivity (for a review see Battaglia-Mayer et al., 2016).

Using the injection site locations with the highest Pearson’s correlation resulted in an overall correlation of the diffusion-based connectivity estimation and of the RLC distribution of r=0.65. This goes in line with the correlation coefficient of r=0.59 reported by Donahue et al. (2016) on connectivity estimated using high resolution DW-MRI tractography and labelled cells counts. These authors studied the predictive power of tractography for connection weights derived from 29 retrograde tracer injections and 91 brain areas, reported by Markov et al. (2014). Although we have used different tractography algorithms and connectivity weights estimation from DW-MRI, both Donahue et al. (2016) and our study show that tractography can indeed estimate structural connectivity weights correlated with the number of measured labelled cells between the cortical areas.

### Tractography misestimated connections

Although tractography produces weighted connectivity proportions showing a good correlation with the proportions of labelled cells, and that most of the weights are in connections with non-zero measured labelled cell count, some connection weights were misestimated. First, across the matching locations and all cortical ROIs, the connection with the most underestimated fraction of diffusion-based connectivity (−0.143) is ROI F2, after injection site in aMIP. This is followed by connection F1-aMIP (−0.135), F1-pMIP (−0.126), F2-aPEip (−0.111) and F2-pPEip (−0.109). Similarly, the most overestimated connectivity from DW-MRI is PGm-aPEip (+0.173), followed by PE-aMIP (+0.151), PE-pMIP (+0.109), S1-aMIP (+0.103) and VIP-aMIP (+0.096). Across the four matching site’s location, tractography misestimated the connectivity the most on ROIs F2, PE, M1, VIP and LIP. The source of these errors can be the intricate white matter geometries and configurations, such as crossing and kissing, causing tractography to follow incorrect orientations (see Jeurissen et al. 2017 and Girard et al., 2020). Additionally, the labelled cells count used in this study was obtained from retrograde axonal tracing, therefore expressing uni-directional connections, while tractography is bi-directional. Thus, asymmetry in the afferent and efferent axon densities of a fascicle could result in a mismatch between the two techniques. Future work should target bi-directional tracing analysis of ROIs with incorrect diffusion-based connectivity estimation.

## MATERIAL AND METHODS

### Neural tracer experiments

#### Subjects, surgical procedures, selection of injection sites, and tracer injections

The tracer experiments were carried out in four male monkeys. In two animals (*Macaca mulatta*; Cases 72 and 73; body weight 12 kg and 12.50 Kg, respectively) retrograde neural tracers were injected at different antero-posterior (A-P) levels of the db-IPS. Additional data from two *Macaca nemestrina* (Cases 10 and 21; body weight 5.2 and 4.4 Kg, respectively), in which a retrograde tracer was injected in the lateral funiculus of the spinal cord, were used for visualizing the origin of corticospinal projections from the db-IPS. Data from these two cases have been already partially used in previous studies (Luppino et al., 1994; Matelli et al., 1998; Rozzi et al., 2006; Borra et al., 2010).

Animal handling as well as surgical and experimental procedures complied with the European law on the humane care and use of laboratory animals (Directives 86/609/EEC, 2003/65/CE, and 2010/63/EU) and Italian laws in force regarding the care and use of laboratory animals (D.L. 116/92 and 26/2014). All procedures were approved by the Veterinarian Animal Care and Use Committee of the University of Rome SAPIENZA or of the University of Parma, and then authorized by the Italian Ministry of Health. Surgical procedures were performed under aseptic conditions. Cases 72 and 73 were pre-anaesthetized with ketamine (5 mg/kg, *i*.*m*.) and dexmedetomidine hydrochloride (0.1 mg/kg; i.m.), intubated and anaesthetized with a mix of Oxygen/ Isoflurane (1-3% to effect). Lidocaine (2%) was used locally to minimize pain during skin incision in the scalp. Desametasone (6mg/kg) was given before dura opening, to prevent brain inflammation and edema. The skull was then trephined over the target region, and the dura was opened to expose the intraparietal sulcus. A constant infusion of Fentanil (0.2mg/kg/h; i.v.) was performed until the end of the surgical procedures. The selection of the injection sites was based on identified anatomical landmarks, such as the rostral tip of the IPS. Once the appropriate site was chosen, fluorescent tracers (Fast Blue [FB] 3% in distilled water, Diamidino Yellow [DY] 2% in 0.2 M phosphate buffer at pH 7.2, both from Dr. Illing Plastics GmbH, Breuberg, Germany) were slowly pressure injected with a glass micropipette attached to the needle of a Hamilton microsyringe at different depths and A-P levels in the medial bank of the IPS. In Case 72, FB (two deposits, 0.15 µl each, at a depth of 3 and 4 mm, respectively) and DY (two deposits, 0.15 µl each, at a depth of 3 and 4 mm, respectively) were injected at about 16 and 13 mm caudal to the rostral end of the left IPS, respectively. In Case 73, FB (0.3 µl) and DY (0.3 µl) were injected at a depth of 4 mm, about 8,5 mm and 18 mm caudal to the rostral end of the right IPS, respectively. After the tracer injections were placed, the dural flap was sutured, the bone was replaced, and the superficial tissues were sutured in layers.

In Cases 10 and 21 in which tracers were injected in the spinal cord, under general anesthesia (Ketamine, 5 mg/kg i.m. and Medetomidine, 0.08–0.1mg/kg i.m.), following a laminectomy, the dura was opened, and the segment of the spinal cord selected for the injection exposed. The retrograde tracer horseradish peroxidase (HRP, 30% in 2% lysolecithin, Sigma-Aldrich, St. Louis, MO) was, then, pressure injected with a 5 µl Hamilton microsyringe in the left lateral funiculus in both monkeys. In one animal (Case 10) the tracer (multiple injections, total amount 10 µl) was injected at the C4-C5 spinal level, in the other (Case 21, multiple injections, total amount 15 µl) at C3--C5 level. Upon the completion of the injections, the spinal cord was covered with Gelfoam and wounds were closed in layers.

During all surgeries, hydration was maintained with saline, and temperature using a heating pad. Heart rate, blood pressure, respiratory depth, O2 saturation, and body temperature were continuously monitored. Upon recovery from anesthesia, the animals were returned to their home cages and closely monitored. Dexamethasone and prophylactic broad-spectrum antibiotics were administered pre- and postoperatively. Furthermore, analgesics were administered intra- and postoperatively.

#### Histological procedures and data analysis

At the end of the survival time (26 days for Case 72; 23 days for Case 73; 3 days for Cases 10 and 21), the animals were given a dose of atropine (0.4 ml; i.m.) and diazepam (Valium, 2ml; i.m.), pre-anaesthetized as above, and received an intravenous lethal injection of sodium thiopental (200 mg/kg; i.v). They were perfused through the left cardiac ventricle with saline, 4% paraformaldehyde, and 5% glycerol in this order. All solutions were prepared in phosphate buffer 0.1 M, pH 7.4. Each brain was then blocked coronally on a stereotaxic apparatus, removed from the skull, photographed, and placed in 10% buffered glycerol for 3 days and 20% buffered glycerol for 4 days. Finally, each brain was cut frozen in coronal sections 60 µm thick. In Cases 10 and 21 the spinal cord was cut in 60 µm thick coronal sections. In Cases 72 and 73, one series of every fifth section was mounted, air-dried, and quickly cover-slipped for fluorescence microscopy. In Cases 10 and 21, one series of every fifth section through the right hemisphere and the brainstem, and every tenth section through the spinal cord was processed for HRP histochemistry using tetramethylbenzidine as the chromogen (Mesulam, 1982). Sections were rinsed in 0.01 M acetate buffer, pH 3.3, and developed at 4°C in a solution of 0.09% sodium nitroferricyanide, 0.005% tetramethylbenzidine, and 0.006% hydrogen peroxide in 0.01 M acetate buffer. Finally, one series of every fifth section in all brains and of every tenth section in the spinal cord in Cases 10 and 21, was stained with the Nissl method (0.1% thionin in 0.1M acetate buffer, pH 3.7).

#### Injection sites and distribution of retrogradely labelled neurons

In Cases 72 and 73, the FB and DY injection sites, defined according to Kuypers and Huisman (1984) and Conde’ (1987), were completely restricted to the cortical gray matter, involving almost the entire cortical thickness, or at least layers III–V. Injection sites were then attributed to area PEip or MIP, as defined from the distribution of corticospinal labelled neurons in the db-IPS (Cases 10 and 21), as detailed in Table 1.

The cortical distribution of FB- and DY-retrogradely labelled cells (Cases 72 and 73) and of HRP-labelled cells (Cases 10 and 21), here referred to as RLC, was plotted in sections every 600 μm (300 μm in Cases 10 and 21), together with the outer and inner cortical borders, using a computer-based charting system. The nomenclature and the map adopted for the areal attribution of the labelled neurons was the same of that used in a recent quantitative study of the connectivity of the parieto-frontal system (Caminiti et al., 2017).

Data from individual sections were then imported into the 3-dimensional (3D) reconstruction software (Demelio et al., 2001) to create volumetric reconstructions of the hemispheres from individual histological sections containing connectional and/or architectonic data and providing realistic visualizations of the labeling distribution. The distribution of RLC on exposed cortical surfaces was visualized in mesial and dorsolateral views of the hemispheres, whereas that in the db-IPS in lateral views of the hemispheres, in which the bank was exposed with dissection of the inferior parietal lobule and the temporal lobe.

#### Quantitative analysis and laminar distribution of retrograde labeling

In all the cases, we counted the number of RLC plotted in the ipsilateral hemisphere, beyond the limits of the injected area, in sections at every 600 μm interval. Cortical afferents to areas PEip or MIP were then expressed in terms of the percentage of labelled neurons found in a given cortical subdivision, with respect to the overall retrograde labeling found for each tracer injection. As for the brain parcellation adopted in this study, for both histological and tractography data, see dedicated paragraph below.

Furthermore, to obtain information about the laminar patterns of the observed connections, the labeling attributed to a given area and reliably observed across different sections and cases was analyzed in sections at every 300 μm in terms of amount of RLC located in the superficial (II–III) versus deep (V–VI) layers.

### Diffusion-weighted MRI experiment

#### Brain processing for ex-vivo DW-MRI acquisition

The diffusion-weighted MRI (DW-MRI) data from ex-vivo brain of a male *Macaca mulatta* (M105, 4 years and 10 months old, 10.1 kg body weight) available from Ambrosen et al. (2020) was used. The brain was perfused following the protocol illustrated in Ahmed et al. (2012) and prepared for MRI ex-vivo scanning as described by Dyrby et al. (2011). The DW-MRI data were acquired at 0.5 mm isotropic resolution. The data were sampled in 180 uniformly distributed directions on each of three b-value shells (b= [1.477, 4.102, 8.040] ms/um^2^) and 9 non-diffusion-weighted images (b=0 ms/um^2^). The protocol was repeated twice and averaged before further processing (for more details on the MRI acquisition protocol, see Ambrosen et al. 2020). We also used the midcortical surface from Ambrosen et al. (2020). The Fiber Orientation Distributions were estimated using the Multi-Shell Multi-Tissue Constrained Spherical Deconvolution algorithm available in the MRtrix3 software (Jeurissen et al., 2014; Tournier et al., 2019). The brain partial volume estimates for the white matter, grey matter, and cerebrospinal fluid were obtained from the averaged non-diffusion-weighted image using the FSL Fast software (Zhang et al., 2001).

#### Brain Parcellation

We used the brain parcellation of the right hemisphere available in Girard et al. (2020). Fifty-nine cortical areas were manually parcellated following the study by Caminiti et al (2017), on the animal used for the ex-vivo DW-MRI acquisition. Areas 46dr and 46dc were grouped in a single regions of interest, (ROI) 46d. Similarly, we grouped areas 46vr, r46vc, c46vc in ROI 46v, areas c12r, i12r, r12r in ROI 12r, areas 9l, 9m in ROI 9, areas 45A, 45B in ROI 45, areas 8Ad, 8Av in ROI 8r&FEF, areas F7PMdr, F7SEF in ROI F7, areas F2vr, F2preCD in ROI F2, areas F5p, F5a/44, F5c in ROI F5. Areas 24 and 25, the insula and Tpt were added to cortical parcellation based on atlases of the rhesus monkey brain (Paxinos et al., 2000; Saleem et al., 2012). Together, these cortical areas make 48 ROIs for investigating the connectivity of PEip and MIP. To obtain a detailed parcellation of the db-IPS, we first merged area PEip and MIP in a single area. This resulted in 38 A-P MRI coronal slices (from #105 to #68; each 0.5 mm thickness) of the db-IPS, then divided into three sectors: dorsal, middle, and ventral. The most anterior part of the area PEip was excluded from the fine parcellation of the db-IPS, because of the difficulty in identifying three sectors. The parcellation was done in the native MRI image space. The MRI images were manually aligned to the stereotaxic plane of the histological sections for visual inspection.

#### DW-MRI Tractography and Connectivity

Probabilistic streamline tractography was performed using the Particle Filtering Tractography algorithms (Girard et al., 2014) available in the DIPY software library (Garyfallidis et al., 2014). Tractography was initiated in all white matter voxels using 25 seeds per voxel (9,713,750 seeds). Streamlines with a length superior to 2 mm in the white matter volume were used as input to the Convex Optimization Modelling for Microstructure Informed Tractography (COMMIT) method (Daducci et al., 2014). COMMIT was used to estimate each streamline contribution (weights) to the intra-axonal MRI signal fraction following the Stick-Zeppelin-Ball white matter microstructure model (Panagiotaki et al., 2012, Daducci et al., 2014). The tractography and microstructure estimation was repeated four times, resulting in a total of 23,137,312 streamlines and weights. All streamlines with an endpoint located in one of the 48 cortical ROIs and an endpoint in the A-P coronal slices of the db-IPS were selected for the diffusion-based connectivity analysis. Streamlines were selected using the MRtrix3 *tck2connectome* (Tournier et al., 2019) command, identifying connected ROIs with a radial search of 1 mm around streamlines endpoints. This resulted in 73,390 streamlines connecting the db-IPS to the cortical areas (dorsal: 29,378; middle: 24,474; ventral: 19,538).

#### Diffusion-based Connectivity Estimation

To cover a similar extent as the tracer injections, we merged the dorsal and middle sectors of the db-IPS. We used a sliding window of 2.5 mm moving in the A-P direction (5 MRI coronal slices) selecting all streamlines connecting the MRI slices to the cortical ROIs. For each sliding window and cortical ROI, we computed the sum of the COMMIT weights (i.e., estimation of the intra-axonal MRI signal fraction) of the streamlines connecting them. The diffusion-based connectivity distribution of a sliding window was obtained by normalizing the sum of the COMMIT weights of each ROI by the sum of COMMIT weights of all ROIs of this sliding window. Thus, for each window located in the db-IPS, the diffusion-based connectivity of a cortical ROI corresponds to its fraction of the COMMIT weights for that window. From the 38 coronal MRI slices (#105 to #68) we obtained 34 (#103 to #70) sliding windows in the A-P extent of the db-IPS, with each window made of 5 consecutive MRI subdivisions (the 2 bordering locations at each extremity of the IPS were excluded). For each sliding window we computed the distribution of diffusion-based connectivity to the selected cortical ROIs. The Pearson’s correlation coefficient was used to compare the diffusion-based connectivity distribution of each window with the histological cell count distributions of the four injection sites.

## Acknowledgments

We acknowledge the support of the Istituto Italiano di Tecnologia to R.C. and the access to the facilities and expertise of the CIBM Center for Biomedical Imaging, founded and supported by Lausanne University Hospital (CHUV), University of Lausanne (UNIL), Ecole Polytechnique Fédérale de Lausanne (EPFL), University of Geneva (UNIGE) and Geneva University Hospitals (HUG). This work was supported by the MIUR of Italy (PRIN 2017 to A.B.M, Grant n. 201794KEER_002 and PRIN 2017 to E.B, Grant n. 2017KZNZLN_002).

## Dedication

This paper is dedicated to our co-author and friend Giorgio M. Innocenti, who passed away unexpectedly on January 12, 2021.

